# Transcriptional fidelity enhances cancer cell line selection in pediatric cancers

**DOI:** 10.1101/2021.10.01.462682

**Authors:** Cuyler Luck, Katharine Yu, Ross A. Okimoto, Marina Sirota

## Abstract

Multi-omic technologies have allowed for comprehensive profiling of patient-derived tumor samples and the cell lines that are intended to model them. Yet, our understanding of how cancer cell lines reflect native pediatric cancers in the age of molecular subclassification remains unclear and represents a clinical unmet need. Here we use Treehouse public data to provide an RNA-seq driven analysis of 799 cancer cell lines, focusing on how well they correlate to 1,655 pediatric tumor samples spanning 12 tumor types. For each tumor type we present a ranked list of the most representative cell lines based on correlation of their transcriptomic profiles to those of the tumor. We found that most (8/12) tumor types best correlated to a cell line of the closest matched disease type. We furthermore showed that inferred molecular subtype differences in medulloblastoma significantly impacted correlation between medulloblastoma tumor samples and cell lines. Our results are available as an interactive web application to help researchers select cancer cell lines that more faithfully recapitulate pediatric cancer.

## Background

Cell lines remain a primary tool for cancer research due to their ease of use, rapid growth, and suitability for a wide variety of experiments. While the merits of their use have been debated for decades, the general consensus is that given careful precautions these models can be good proxies for their tumors of origin^1,2^. In the -omics age, a large number of cancer cell lines have been comprehensively characterized based on their genetics^3–6^, drug sensitivities^3,4^, transcriptomics^3–6^, proteomics^5,7^, metabolomics^5,8^, and genetic dependencies^9,10^. The wealth of annotations for these models has supported their use in understanding cancer and will continue to drive innovation in the future.

In 2021, over 11,000 children and adolescents in the US will be diagnosed with cancer and over 1,200 will die of their disease^11^. While survival rates have dramatically improved for pediatric cancer since the 1970s, incidence has been slightly increasing for unknown reasons and roughly 15% of patients will succumb to their illness^11^. For the many who now survive childhood cancer, the impact of therapy-related toxicities remains a significant challenge^12,13^. Pediatric cancers are thought to be biologically distinct from adult cancers in multiple ways^14–16^, and as such it is worth considering whether the same cell lines that effectively represent adult tumors will also reflect pediatric cancers.

Choosing the most optimal cell lines that faithfully recapitulate a specific tumor type can potentially enhance clinical translation. Previous work from our lab has leveraged massive publicly available transcriptomic datasets from The Cancer Genome Atlas (TCGA) and the Cancer Cell Line Encyclopedia (CCLE) to create a resource that enables researchers to identify the most representative cell lines for a specific tumor type of interest^17^. This prior analysis largely focused on adult tumor types. Similarly, a more recent study using both adult and pediatric samples developed an unsupervised alignment method capable of resolving intrinsic cell line – tumor differences, revealing that the majority of cell lines align with their tumor types of origin with higher transcriptional similarity than previously thought^18^. Moreover, through the development of a prototype pediatric cancer dependency map the same method was employed to show that 74% of pediatric cell lines had expression patterns that matched those of their primary tumors, and that cell lines derived from pediatric cancers generally cluster well with their primary sample counterparts^10^. While informative, these studies largely focused on adult tumor types and did not explore specific molecular subtypes in the context of pediatric tumors.

In this study we will focus on how pediatric cancer cell lines reflect their native tumors and demonstrate how transcriptional subclassification within certain tumor types can better inform cell line selection. Specifically, we use a transcriptomics-based comparison between 1,655 pediatric tumor samples and 799 cell lines to determine the most representative cell lines for 12 pediatric cancers. These results are available as an interactive web application (https://pecanexplorer.org/) to help researchers choose the most representative cell lines for their needs. Furthermore, through our transcriptional analysis of medulloblastomas, we highlight the importance of molecularly informed selection of cell lines to more accurately reflect subtype specific cancers. Collectively, we provide a user-friendly and interactive platform to inform and guide the cancer community in the selection of cell lines that more faithfully recapitulate patient-derived pediatric cancers.

## Results

### Global transcriptomic analysis of pediatric tumor samples and cell lines demonstrates biologically-meaningful relationships within and between diseases

To choose pediatric tumor samples, we used the Treehouse Tumor Compendium v11 Public PolyA dataset and filtered for samples with an age at diagnosis less than or equal to 18 years. We further subset the data to diseases with 30 or more tumor samples, after including glioblastoma multiforme and gliomatosis cereberi as “glioma” samples. For cell line selection, we used Treehouse’s Cell Line Compendium v2 and retained cell lines which had TCGA codes defined in the Broad Institute’s CCLE annotations file. Cell lines were not filtered based on patient age. Together, this yielded 1,655 tumor samples and 799 cell lines for analysis (Figure 1a, Tables 1 and 2).

**Table 1.** Pediatric tumor samples. Distribution of 1,655 pediatric tumor samples among 12 tumor types.

| Tumor Disease | Num. Samples | Tumor Disease | Num. Samples |
| --- | --- | --- | --- |
| Acute lymphoblastic leukemia | 402 | Glioma | 197 |
| Acute myeloid leukemia | 241 | Medulloblastoma | 96 |
| Alveolar rhabdomyosarcoma | 43 | Neuroblastoma | 195 |
| Embryonal rhabdomyosarcoma | 53 | Rhabdomyosarcoma | 44 |
| Ependymoma | 67 | Osteosarcoma | 117 |
| Ewing sarcoma | 65 | Wilms Tumor | 135 |

**Table 2.** Cancer cell lines. Distribution of 799 cancer cell lines among 29 TCGA codes.

| TCGA Code | Num. Lines | TCGA Code | Num. Lines | TCGA Code | Num. Lines |
| --- | --- | --- | --- | --- | --- |
| ALL | 30 | LAML | 31 | OV | 41 |
| BLCA | 22 | LCML | 14 | PAAD | 39 |
| BRCA | 48 | LGG | 9 | PRAD | 7 |
| CLL | 4 | LIHC | 24 | SARC | 29 |
| COAD/READ | 49 | LUAD | 70 | SCLC | 49 |
| DLBC | 36 | LUSC | 22 | SKCM | 45 |
| ESCA | 22 | MB | 4 | STAD | 36 |
| GBM | 31 | MESO | 9 | THCA | 10 |
| HNSC | 29 | MM | 25 | UCEC | 28 |
| KIRC | 21 | NB | 15 |  |  |

**Fig. 1.**
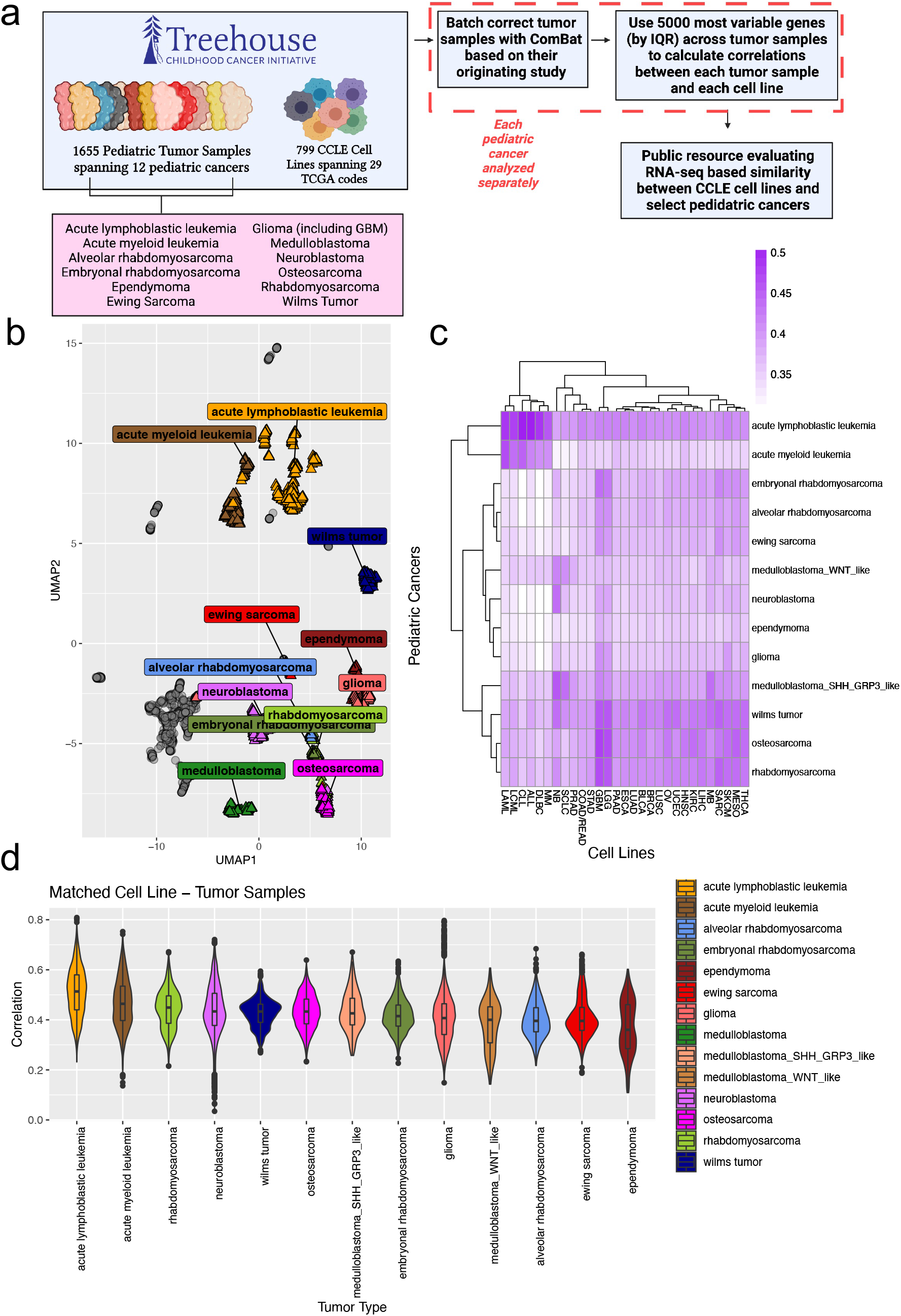
Global overview of transcriptomic comparison between pediatric tumor samples and cancer cell lines. (a) Graphical visualization of input data and processing performed for web application, created with BioRender.com. (b) UMAP of all cell line and tumor samples together using TPM data, based on the top 5,000 most variable genes by IQR across all samples. Cell lines are represented as gray circles, tumor samples are represented as colored triangles. (c) Heatmap of mean Spearman correlation values summarized to cell line TCGA code and tumor sample disease levels, correlation values displayed were aggregated after individual calculation for each tumor sample disease. (d) Violin plot of Spearman correlation values between pediatric tumor samples and the closest manually matched cell line disease type, data aggregated after individual calculation for each tumor sample disease.

Using our curated pediatric dataset, we performed dimensionality reduction using UMAP to visualize all tumor samples and cell lines based on the 5000 most variable genes (measured by IQR) across all samples. We observed separation between tumor samples and cancer cell lines on UMAP 1, while blood versus solid cancers separate out on UMAP 2 (Figure 1b). Strikingly, even such a simple analysis is sufficient to cluster tumor samples by disease that are organized in rather biologically sensible ways. With these findings, we hypothesized that dividing the data into tumor samples and cell lines before generating similar UMAPs would allow for even more precise clustering of samples by disease type. Indeed, we observed that blood (leukemia, lymphoma, myeloma) cancer cell lines appeared to be more transcriptionally similar, suggesting a less heterogeneous and more clonal origin when compared to their solid tumor counterparts (Supplemental Figure 1a and 1b).

To quantify similarity between tumor samples and cell lines using TPM data, we batch corrected tumor data within each disease type using ComBat^19,20^ to remove study-specific differences (Supplemental Figures 2-6). We then calculated Spearman correlation coefficients between tumor samples of a given disease type and all cell lines based on the 5000 most variable genes across the disease-specific tumor samples. Such an approach allows us to use correlation coefficients as measures of how well a given cell line transcriptionally represents the majority of tumor samples within a disease.

A global view of mean correlations summarized by cell line TCGA code and tumor sample disease revealed patterns which would be expected based on their underlying biology (Figure 1c). Blood cancers form a distinct cluster from solid tumors for both tumor samples and cell lines and correlate the best with other blood cancers rather than with solid tumors. With the exception of Sonic Hedgehog (SHH)/Group 3-like medulloblastoma, four of the five central nervous system tumor types form a cluster adjacent to a smaller cluster of Ewing sarcoma and embryonal and alveolar rhabdomyosarcomas. Tumor types that had similarly high correlations with multiple cell line diseases form a cluster towards the bottom of the heatmap, comprised of SHH/Group 3-like medulloblastoma, Wilms Tumor, osteosarcoma, and rhabdomyosarcoma with no subtype information.

Interestingly, specific sarcoma tumor types (Ewing sarcoma, rhabdomyosarcoma) had particularly high mean correlations with cell lines representing glioblastoma multiforme (GBM) or low grade glioma (LGG). This is in line with the original Celligner analysis, which found that almost all cell lines representing the central nervous system were classified as soft tissue tumors using Celligner-aligned data^18^. Further rationale for these observations include the upregulation of neural crest gene signatures in small round cell tumors, including Ewing sarcoma family tumors, which underly its overlapping classification with peripheral Primitive Neuroectodermal Tumors (pPNET)^21^.

We were also interested in evaluating how well tumor samples correlated to cell lines whose diseases should be similarly aligned. After manually determining such pairings (see Methods), we observed that the two pediatric blood cancers (AML and ALL) best correlated with their matched cell lines, with acute lymphoblastic leukemia samples having a notably higher median correlation than any other tumor type (Figure 1d).

Correlation coefficients calculated in our analysis are lower than those reported when Celligner is implemented^18^, likely reflecting intrinsic tumor sample – cell line differences that deflate such values. Consequently, we caution against directly comparing values generated in this study to those determined after implementing unsupervised alignment methods.

### Identifying best-fit cell lines for pediatric cancers reveals that most tumor types are best reflected by a corresponding cell line disease

Of the tumor types we analyzed, 8/12 (66%) correlated the most highly with a cell line derived from the most similar disease type available. Within those that did not match their most similar disease type, we did not have any Wilms Tumor (WT) cell lines in our data and WT tumor samples were instead expected to best correlate with kidney renal clear cell carcinoma (KIRC) cell lines. However, the best correlated cell line for WT tumor samples was SNU-182, a hepatocellular carcinoma cell line. Ependymoma tumor samples correlated best with glioblastoma multiforme (GBM) cell lines, with its top 6 highest correlation cell lines being GBM. This was somewhat surprising as we expected these samples to better correlate with low grade glioma (LGG) cell lines, the best performing of which was the ninth most well correlated cell line with ependymoma tumor samples. However, it is possible that the ependymoma tumor samples were higher grade than we expected and consequently reflect GBM cell lines better than LGG cell lines. Medulloblastoma tumor samples correlated best with small cell lung cancer (SCLC, 7 of top 10 correlated cell lines) and neuroblastoma (NB, 3 of top 10 correlated cell lines) cell lines, while osteosarcoma tumor samples correlated best with GBM cell lines (7 of top 10 correlated cell lines) although one sarcoma cell line was present in the top 10 cell lines for this disease (Hs 729.T, an embryonal rhabdomyosarcoma cell line).

To demonstrate the results of our analysis we focused on two case studies: one solid tumor (Ewing sarcoma, ES) and one blood cancer (acute myeloid leukemia, AML). Ewing sarcoma tumor samples had almost no visible batch effects present prior to ComBat correction (Supplemental Figure 3c), and consequently such an adjustment was not as profound as with other tumor types. When aggregated by TCGA code, the cell line diseases with the highest median correlations with ES tumor samples were GBM and LGG cell lines (Figure 2a). However, sarcoma (SARC) cell lines had the third highest median correlation and had the highest maximum correlations of any cell line disease type. When focusing on the top 10 cell lines in terms of median correlation with ES tumor samples, the top six performers were all SARC cell lines and all six of these were ES cell lines (5 Ewing sarcoma, 1 Askin tumor) (Figure 2b). Only one other ES cell line exists in our dataset, CADO-ES-1, which was the 11^th^-highest cell line in terms of median correlation and interestingly has an EWSR1-ERG fusion while the other six ES cell lines all possess EWSR1-FLI1 fusions. This group of EWSR1-FLI1 ES cell lines preceded four GBM cell lines in the top 10 performers which were notably less well correlated. When looking at correlations between ES tumor samples and SARC cell lines, the EWSR1-FLI1 SARC cell lines which best correlated with ES tumor samples form a distinct cluster from other SARC cell lines (Supplemental Figure 8b) and clearly better reflect a larger proportion of ES tumor samples than do cell lines in the other cluster. Notably, CADO-ES-1 displays a pattern of correlations similar to the EWSR1-FLI1 cell lines, but was assigned to a different cluster than the EWSR1-FLI1 cell lines.

**Fig. 2.**
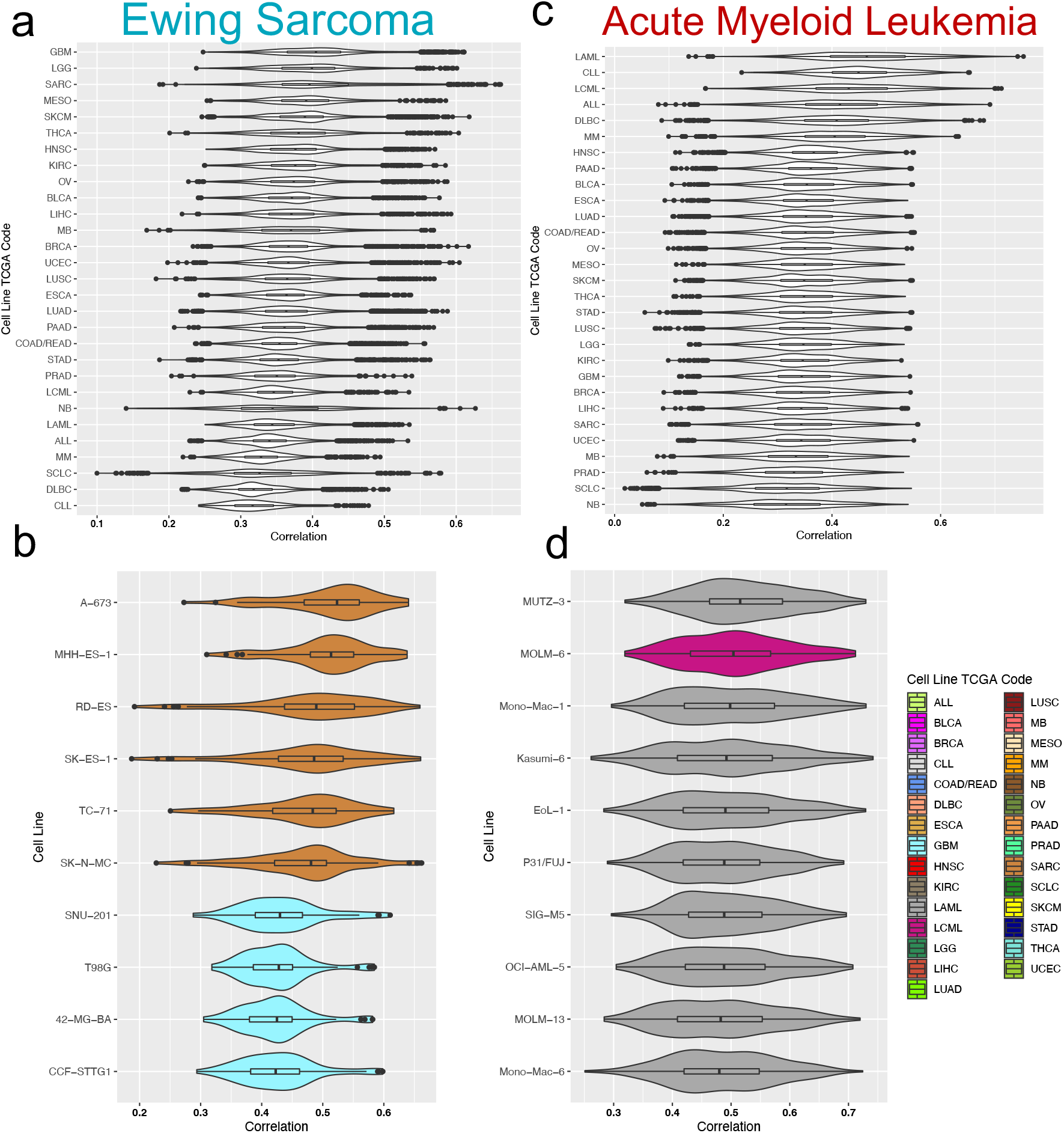
Case study analyses for Ewing sarcoma and acute myeloid leukemia. (a) Violin plot of Spearman correlation values between Ewing sarcoma tumor samples and all cancer cell lines, grouped by cancer cell line TCGA code. (b) Violin plot of Spearman correlation values between Ewing sarcoma tumor samples and the top 10 cancer cell lines based on median correlation value. (c) Violin plot of Spearman correlation values between acute myeloid leukemia tumor samples and all cancer cell lines, grouped by cancer cell line TCGA code. (d) Violin plot of Spearman correlation values between acute myeloid leukemia tumor samples and the top 10 cancer cell lines based on median correlation value.

AML tumor samples displayed a clear batch effect based on the study the samples originated from which was effectively removed after ComBat adjustment (Supplemental Figure 2b). The six blood cancer cell line groups (LAML, CLL, LCML, ALL, DLBC, and MM) had substantially higher median correlations with the AML tumor samples than any solid tumor cell line diseases (Figure 2c). Furthermore, AML cell lines (TCGA code LAML) had the highest median correlation with AML tumor samples, suggesting that these cell lines do indeed represent pediatric AML tumor samples the best of any blood cancer cell line diseases. Interestingly, of the top 10 individual cell lines based on median correlations, 9/10 were AML cell lines while a BCR-ABL positive LCML cell line (MOLM-6) was the second-best performer. Within AML tumor sample versus cell line correlations, four general clusters of cell lines are apparent (Supplementary Figure 7b).

PCA plots (pre- and post-ComBat correction), violin plots summarized by cell line disease, top 10 performing cell lines, and matched-disease heatmaps are all available for each tumor disease in the accompanying interactive R Shiny app for this publication at https://pecanexplorer.org/.

### Inferred subtypes in medulloblastoma samples alter cell line comparisons

When we initially analyzed the 96 pediatric medulloblastoma tumor samples, we noticed that ComBat correction did not solve apparent batch effects in our data (Supplementary Figure 11 a and b). Since medulloblastoma is a disease comprised of four distinct molecular subtypes^22,23^, we suspected that what appeared to be study-specific batch effects may in fact be distinct transcriptional signatures arising from subtype differences. We plotted log-transformed expected counts data for the medulloblastoma samples and found that they formed two clear groups along the first principal component (Figure 3a), notably with some samples from the same study belonging to both groups, further suggesting this was not simply a study-specific effect. We performed K-means clustering using k=2 to identify two groups along PC1 which form distinct clusters (Figure 3b).

**Fig. 3.**
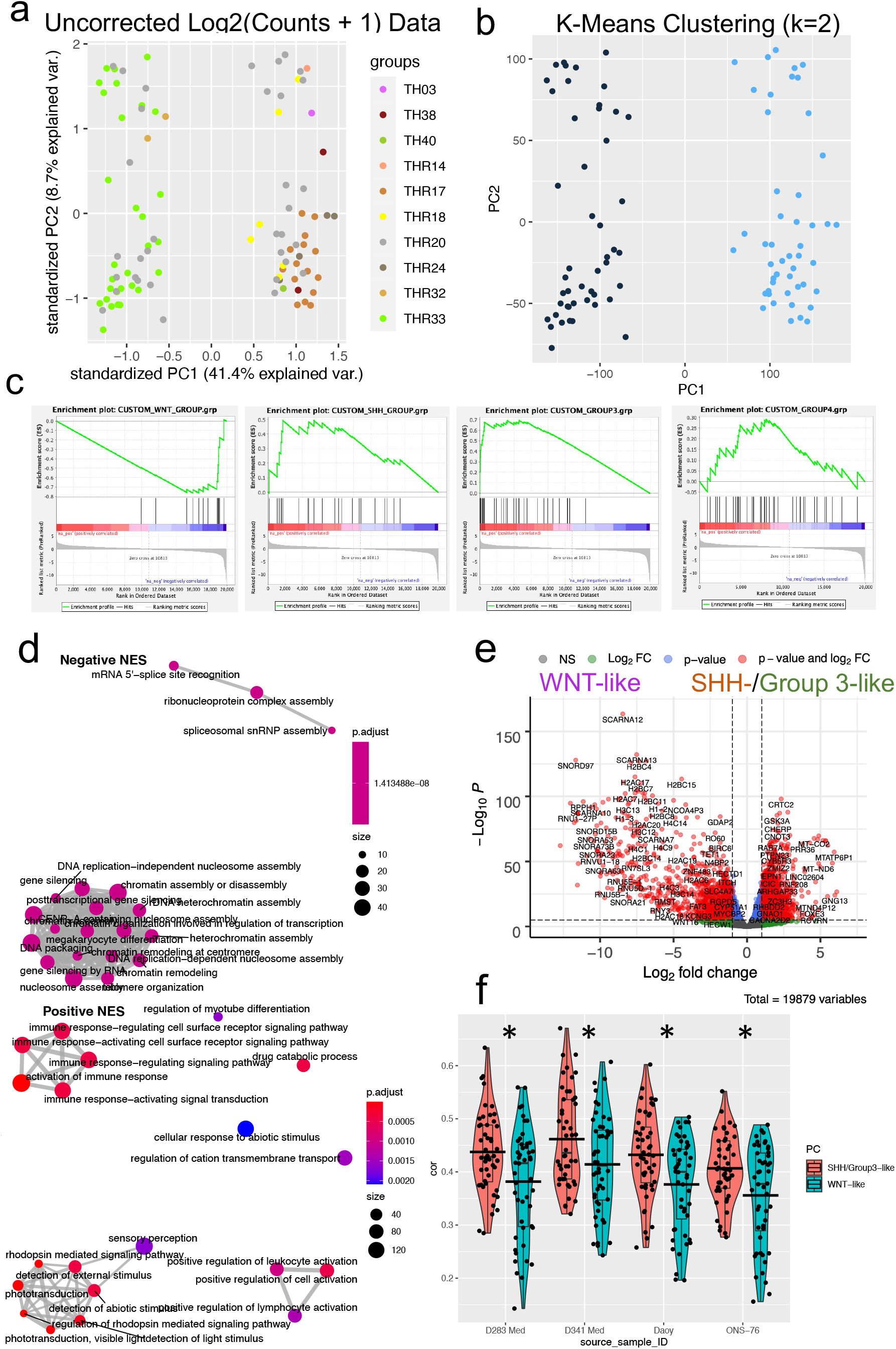
Inferred medulloblastoma tumor sample molecular subtypes and their impact on correlations with medulloblastoma cell lines. (a) PCA plot of log_2_-transformed medulloblastoma tumor sample expected counts data using the top 5000 most variable post-transformation genes by IQR. (b) K-means clustering (k=2) of PCA plot from (a) reveals two clear groups divided along principal component 1. (c) GSEAPreranked analysis of edgeR output comparing the two groups from (b) using custom gene sets for each of the four medulloblastoma molecular subtypes. (d) Unbiased GSEA analysis of edgeR output comparing the two groups from (b) using the gseGO function from clusterProfiler. (e) EnhancedVolcano plot of edgeR output comparing the two groups from (b). (f) Spearman correlation values between medulloblastoma tumor samples and the four medulloblastoma cell lines, separated by inferred molecular subtype of the tumor samples. Asterisk represents FDR adjusted p value < 0.05, crossbars indicate mean values.

The clinical data available from Treehouse did not specify subtype-level diagnoses for medulloblastoma tumor samples, so we chose to use differential expression analysis to infer subtypes from the available expression data. Using expected counts data we ran a simple edgeR^24–28^ comparison between the two groups identified by k-means clustering. We then leveraged prior work from Hooper et al.^22^ who previously found gene sets associated with each molecular subtype of medulloblastoma. We used the contributing genes in these gene sets to perform Gene Set Enrichment Analysis (GSEA) for each subtype (Figure 3c) and found that WNT-related genes were strongly downregulated (FDR q = 0.0021) in the data (higher in the cluster with a negative PC1), while SHH-related and Group 3-related genes were strongly upregulated (FDR q = 0.059, FDR q = 0.00, respectively). Group 4-related genes were not significantly up or downregulated (FDR q = 0.42). Together, these results suggested that WNT tumor samples and SHH/Group 3 tumor samples may be driving the observed clustering patterns.

To ask what pathways were significantly up or downregulated in our edgeR output without biasing the gene sets, we used gene ontology analysis (Figure 3d). Nucleosome assembly and RNP assembly terms were highly significant in the group associated with expression of WNT-related genes. Immune response and phototransduction signaling pathway terms were significant in the group associated with SHH/Group 3-related genes. This further aligns with Hooper et al.’s work showing that the SHH gene cluster was relatively related to immune-response pathways, while the Group 3 gene cluster was closely related to rod development pathways^22^. Thus, an unbiased query of pathways associated with each sample cluster further supports the notion that WNT and SHH/Group-3 samples define the two groups. At the gene level, a volcano plot of the data reveals that a large number of genes are significantly differentially expressed between the two groups, including many small nucleolar RNAs, small Cajal body-specific RNAs, and histone-associated proteins that are present at a significantly higher level in the WNT-like group (Figure 3e).

Since these two groups defined by WNT-like and SHH/Group 3-like samples have clearly different transcriptomic signatures, we wondered whether this would impact how well they are modeled by medulloblastoma cell lines. There are four medulloblastoma cell lines in our analysis, for which some lineage and molecular subtype information is available (D283 MED - Group 3/4, MYC Exp; D341 - Group 3, MYC Amp; DAOY – SHH, non MYC; and ONS76 – SHH, non MYC)^32,33^. We aggregated medulloblastoma cell line – medulloblastoma tumor sample correlations based on whether the tumor samples belonged to the WNT-like or SHH/Group 3-like group and compared the two groups at a bulk level (Supplemental Figure 11c) as well as at the individual medulloblastoma cell line level (Figure 3f). Strikingly, the SHH/Group 3-like tumor samples had significantly higher mean correlations than the WNT-like tumor samples when aggregated (Wilcoxon p-value = 2.2e-06) and across all four cell lines (Wilcoxon FDR adjusted p-values: D283 MED – 0.024, D341 – 0.024, ONS-76 – 0.024, DAOY – 0.018). This makes biological sense given the subtype information available for the four medulloblastoma cell lines in this analysis, and yet it underscores a key point for this disease – medulloblastoma cell lines do not represent all medulloblastoma tumor samples equally.

Given our success in inferring molecular subtypes in medulloblastoma, we performed a similar analysis in two other tumor types thought to have subtypes: glioblastoma and neuroblastoma. GBM was originally suggested to be divisible into four subtypes^36^, which has since been revised to three subtypes^37^. We attempted to use a 500-gene set recently used with TCGA samples^37^ to classify our glioma samples and found that using 474 of the genes that were both in our data and the 500-gene set did not appear to clearly enhance clustering compared to using the top 5000 most variable genes (by IQR) across the tumor samples (Supplemental Figure 12 a, b, and c). Subsetting our data to just GBM tumor samples and applying the same gene set did not seem to define any clear groupings (Supplemental Figure 12 d and e). Neuroblastoma has also been recently divided into four epigenetic subtypes driven by super-enhancers^38^. We used the list of target genes for the defined super-enhancers to group our neuroblastoma tumor samples into subtype groups, but saw little difference in clustering compared to using the top 5000 most variable genes (by IQR) across tumor samples (Supplemental Figure 13).

## Discussion

Despite their limitations, cell lines continue to be the primary drivers of cancer research. With an ever-growing wealth of annotations, it continues to become simpler for researchers to choose cell lines that best model their tumor type of interest. Here we have provided an interactive resource for those studying pediatric cancers to evaluate how well cell lines reflect any of 12 pediatric tumor types (https://pecanexplorer.org). We contribute this work as an additional tool to use in conjunction with previous work from our lab and others, as many different characteristics of cell lines may impact their suitability as models for a particular project.

We found that pediatric cancers generally are well represented by cell lines, as 8/12 (66%) of diseases had a cell line of the matched disease type as the best correlated cell line. For those diseases that were not well represented, particularly medulloblastoma and osteosarcoma, these tumor types may be difficult to transcriptionally mimic in culture and more reflective models are needed. For osteosarcoma, a tumor type rife with diversity in genomic alterations, choosing a cell line may instead require a larger focus on whether the structural rearrangements and mutations present in a cell line match the research question at hand. When analyzing Wilms Tumor, we had no WT cell lines to directly compare the tumor samples with. A number of WT cells have recently been described^34^, and should be harmonized with existing data so researchers can evaluate for themselves how well these lines model primary tumor samples.

As we have shown here, it may be important to consider molecular subtype when evaluating how well cell lines reflect a disease of interest. This is particularly of concern in medulloblastoma, where the vast majority of cell lines available are derived from SHH and Group 3 tumors, or simply lack any subtype information^32^. As previously noted, there are very few WNT and Group 4 cell lines available (1 and 2 respectively as of 2016^32^), which is troubling as around half of diagnoses are classified as one of these subtypes. Indeed, the last time that a literature search was performed it was found that the four medulloblastoma cell lines used in this work accounted for over 72% of raw citations among medulloblastoma lines^32,35^. In light of our results showing that these cell lines differentially correlate with medulloblastoma tumor samples with different inferred molecular subtypes, we add to the call for the development of newer cell lines that better represent WNT and Group 4 medulloblastoma.

Our study has limitations that should be considered when comparing results with others’ work. We lack information on tumor purity which we have previously shown can confound correlations between cell lines and tumor samples in many cancer types^17^, and as a result we may be underestimating correlation coefficients. Similarly, as we did not computationally account for intrinsic differences between tumor samples and cell lines via a method such as Celligner^10,18^, this could further underestimate absolute correlation coefficient values. However, this should only impact study – study comparisons of correlation coefficient values and should not alter our results when compared within this work. We also note that our work only reflects transcriptomic similarity between tumor samples and cell lines, which may not be the most important factor in choosing suitable cell lines for a research project. Researchers should be sure to carefully consider the underlying biology that their project investigates, for example by paying close attention to structural rearrangements in osteosarcoma or acute lymphoblastic leukemia. With this in mind, we hope that our work can add another point of view to be considered when choosing cell lines to reflect pediatric cancer *in vitro*.

As more comprehensive molecular information becomes available for patient samples and cell lines alike, the activation energy for using such information to drive project design becomes lower. We hope that future work generating cell lines for pediatric cancers which currently have poor models will lead to more precise research with more positive results. Given the impact that molecular subtypes may have on choosing an *in vitro* model, we also call for the continued comprehensive profiling of tumor samples that are deposited in public compendia. As our models improve at reflecting true disease, so too will our understanding of tumor biology and therapies.

## Methods

### Data selection

To avoid any artificial differences arising from the use of different computational processing approaches, we took advantage of the University of California, Santa Cruz Treehouse Childhood Cancer Initiative’s publicly available repositories which use the Toil pipeline to unify RNA-seq samples from different studies. For tumor sample data we used the Tumor Compendium v11 Public PolyA (https://treehousegenomics.soe.ucsc.edu/public-data/#tumor_v11_polyA), which has log2(TPM+1) and expected count expression data available for 12,747 samples from TARGET, TCGA, Treehouse, and other studies. For cell line data we used the Cell Line Compendium v2 (https://treehousegenomics.soe.ucsc.edu/public-data/previous-compendia.html#cell_line_v2)., which has log2(TPM+1) expression data available for 912 cell lines from CCLE, Treehouse, and other repositories.

To focus on pediatric cancers, we used the clinical information file from Treehouse’s Tumor Compendium v11 Public PolyA to subset the tumor samples to those where the patient’s age at diagnosis was equal to or less than 18 years. This filtering resulted in 1,893 samples spanning 83 unique disease types, although the vast majority of diseases consisted of a low number of samples (n = 71 diseases with low samples). To ensure adequate sample size for sufficient statistical analyses we chose to further restrict the data to tumor types with 30 or more samples, which resulted in 1,626 samples across 12 pediatric cancers. We also chose to include glioblastoma multiforme samples (GBM, n=29) and gliomatosis cereberi (GC, n=1) within the broader classification of glioma samples (n=168 before addition of GBM and GC) as these diseases are subtypes of gliomas but did not have enough samples to be analyzed on their own. After the addition of GBM and GC samples, we had a final cohort of 1,655 pediatric tumor samples spanning 12 pediatric cancers.

For cell line data, we used Treehouse’s Cell Line Compendium v2. In order to obtain the highest-confidence cell line disease information, we used the publicly available Cancer Cell Line Encyclopedia (CCLE) annotation file (https://depmap.org/portal/download/, “Cell_lines_annotations_20181226.txt”) to define cell line diseases based on the The Cancer Genome Atlas (TCGA) code each cell line was classified under (using the tcga_code variable). This stratified the cell lines in our dataset to CCLE lines with TCGA code information available, resulting in 799 cell lines available for use in our study. Most TCGA code abbreviations are explained at https://gdc.cancer.gov/resources-tcga-users/tcga-code-tables/tcga-study-abbreviations. For those that are not, ALL = acute lymphoblastic leukemia, CLL = chronic lymphocytic leukemia, MB = medulloblastoma, MM = multiple myeloma, NB = neuroblastoma, and SCLC = small cell lung cancer.

### Tumor sample batch correction and correlation analysis

We first removed all genes with zero variance from the cell line data. We cycled through all tumor types, first using ComBat^19,20^ to batch correct samples based on the site ID encoded in each tumor sample’s Treehouse sample ID. We then removed genes with no variance from the tumor data, and generated PCA plots using the top 5000 most variable genes by IQR pre- and post-batch correction. Next, we inner joined the cell line and disease-specific tumor data together such that the only genes in the combined data were those with non-zero variance in both datasets. Finally, we subset the merged data to the top 5000 most variable genes by IQR across the tumor samples only and used the output to calculate Spearman correlation coefficients between all cell lines and every tumor sample in a given disease.

For matched disease comparisons between tumor samples and cell lines, the following disease – TCGA code pairings were used (Tumor disease – cell line TCGA Code): Medulloblastoma (either inferred subtype group) – MB; Ewing sarcoma – SARC; acute lymphoblastic leukemia – ALL; alveolar rhabdomyosarcoma – SARC; osteosarcoma – SARC; embryonal rhabdomyosarcoma – SARC; glioma – LGG and GBM; acute myeloid leukemia – LAML; Wilms tumor – KIRC; neuroblastoma – NB; ependymoma – LGG; and rhabdomyosarcoma – SARC.

### Differential expression and GSEA

For differential expression (DE) analysis of medulloblastoma tumor samples we used expected count expression data and defined two groups of samples based on whether they had a positive or negative principal component 1 in the PCA plot shown in Figure 3a. We used a standard edgeR pipeline per the user’s manual for DE analysis, with the exactTest and p.adjust functions used to produce p-values and FDR adjusted p-values. Then, we converted gene IDs (originally in Ensembl ID format) to Hugo gene symbols using biomaRt^39,40^, conservatively removing genes where duplicate Ensembl ID or Hugo symbols were generated during conversion. We used the EnhancedVolcano^41^ function to generate the volcano plot. For GSEA we took Hugo-named fold changes from the edgeR output and used them as input into the gseGO function from clusterProfiler^31^, visualizing the data with emapplot^42^. For GSEAPreranked^29,30^ we used the desktop version of the Broad Institute’s application (available at http://www.gsea-msigdb.org/gsea/index.jsp) where the input was the Hugo-named fold changes from edgeR and the custom gene sets were defined as explained in the main text. Non-default settings were the use of no collapsing to remap gene symbols, and a minimum size of 10 to accommodate the small custom WNT gene set.

## Data Availability

All data used in this study are publicly available. Treehouse tumor and cell line data are available at https://treehousegenomics.soe.ucsc.edu/public-data/. Tumor TPM and expected counts data were from the Tumor Compendium v11 Public PolyA (April 2020). Cell line TPM data were from the Cell Line Compendium v2 (December 2019). CCLE cell line annotations can be downloaded at https://depmap.org/portal/download/ (“Cell_lines_annotations_20181226.txt”).

## Code Availability

The source code for data selection and analysis are available at https://github.com/cuylerluck/treehouse_pediatric_paper with a guide for reproducible use.

## Acknowledgements

The authors thank the members of the Sirota and Okimoto labs for their valuable insight. The authors further thank Dr. Tomiko Oskotsky and Dr. Boris Oskotsky for their help in publishing the R shiny application. This work was in part supported by a NIGMS Predoctoral Training in Biomedical Sciences T32 (C.L., T32 GM 136547), an NCI R37 Merit Award (R.A.O., R37CA255453), and an NIH-NCI Career Development Award (R.A.O., K08CA222625).

## Author Contributions

Conceptualization, M.S., C.L., K.Y.; investigation, C.L., K.Y.; software, C.L.; writing of the original draft, C.L.; discussion, C.L., K.Y., R.A.O., M.S.; writing (review and editing), C.L., K.Y., R.A.O., M.S.; supervision R.A.O. and M.S.

## Author Information

Department of Medicine, University of California, San Francisco, San Francisco, California, USA

Cuyler Luck, Ross A. Okimoto

Bakar Computational Health Sciences Institute, University of California San Francisco, San Francisco California, USA

Cuyler Luck, Katharine Yu, Marina Sirota

Department of Pediatrics, University of California, San Francisco, San Francisco California, USA.

Cuyler Luck, Katharine Yu, Marina Sirota

Helen Diller Family Comprehensive Cancer Center, University of California, San Francisco, San Francisco, California, USA

Ross A. Okimoto

## Supplementary Information

Accompanies this paper as a separate file.

## Supplementary Figure Legends

**Supp. Fig. 1. Individual dimensionality reductions of tumor samples and cancer cell lines**.

(a) UMAP of pediatric tumor samples using TPM data prior to any further analysis, using the top 5000 most variable genes by IQR across all tumor samples. (b) UMAP of cancer cell lines using TPM data prior to any further analysis, using the top 5000 most variable genes by IQR across all cancer cell lines.

**Supp. Fig. 2. ComBat batch correction of acute lymphoblastic leukemia, acute myeloid leukemia, and alveolar rhabdomyosarcoma tumor samples**.

Pre- (left) and post- (right) ComBat batch corrected PCA plots using the top 5000 most variable genes by IQR for acute lymphoblastic leukemia (a), acute myeloid leukemia (b), and alveolar rhabdomyosarcoma (c).

**Supp. Fig. 3. ComBat batch correction of embryonal rhabdomyosarcoma, ependymoma, and Ewing sarcoma tumor samples**.

Pre- (left) and post- (right) ComBat batch corrected PCA plots using the top 5000 most variable genes by IQR for embryonal rhabdomyosarcoma (a), ependymoma (b), and Ewing sarcoma (c).

**Supp. Fig. 4. ComBat batch correction of glioma, SHH-/Group 3-like medulloblastoma, and WNT-like medulloblastoma tumor samples**.

Pre- (left) and post- (right) ComBat batch corrected PCA plots using the top 5000 most variable genes by IQR for glioma (a), SHH-/Group 3-like medulloblastoma (b), and WNT-like medulloblastoma (c).

**Supp. Fig. 5. ComBat batch correction of neuroblastoma, osteosarcoma, and rhabdomyosarcoma tumor samples**.

Pre- (left) and post- (right) ComBat batch corrected PCA plots using the top 5000 most variable genes by IQR for neuroblastoma (a), osteosarcoma (b), and rhabdomyosarcoma (c).

**Supp. Fig. 6. ComBat batch correction of Wilms tumor samples**.

Pre- (left) and post- (right) ComBat batch corrected PCA plots using the top 5000 most variable genes by IQR for Wilms tumor.

**Supp. Fig. 7. Heatmaps of disease-specific matched correlations for acute lymphoblastic leukemia, acute myeloid leukemia, alveolar rhabdomyosarcoma, and embryonal rhabdomyosarcoma**.

Heatmaps of correlations between tumor samples and manually matched cell lines (based on the most similar available cell line TCGA code for a given disease) are shown for acute lymphoblastic leukemia (a), acute myeloid leukemia (b), alveolar rhabdomyosarcoma (c), and embryonal rhabdomyosarcoma (d).

**Supp. Fig. 8. Heatmaps of disease-specific matched correlations for ependymoma, Ewing sarcoma, glioma, and SHH- /Group 3-like medulloblastoma**.

Heatmaps of correlations between tumor samples and manually matched cell lines (based on the most similar available cell line TCGA code for a given disease) are shown for ependymoma (a), Ewing sarcoma (b), glioma (c), and SHH- /Group 3-like medulloblastoma (d).

**Supp. Fig. 9. Heatmaps of disease-specific matched correlations for WNT-like medulloblastoma, neuroblastoma, osteosarcoma, and rhabdomyosarcoma**.

Heatmaps of correlations between tumor samples and manually matched cell lines (based on the most similar available cell line TCGA code for a given disease) are shown for WNT-like medulloblastoma (a), neuroblastoma (b), osteosarcoma (c), and rhabdomyosarcoma (d).

**Supp. Fig. 10. Heatmap of disease-specific matched correlations for Wilms tumor**.

Heatmap of correlations between tumor samples and manually matched cell lines (based on the most similar available cell line TCGA code for a given disease) is shown for Wilms tumor.

**Supp. Fig. 11. Medulloblastoma tumor sample TPM batch correction and bulk correlation comparisons broken down by tumor sample inferred subtype**.

(a) Pre-ComBat batch correction PCA of medulloblastoma tumor samples based on the top 5000 most variable genes by IQR across all tumor samples. (b) Post-ComBat batch correction (based on sample site) PCA of medulloblastoma tumor samples based on the top 5000 most variable genes by IQR across all tumor samples. (c) Violin plot of Spearman correlation values between medulloblastoma cell lines and medulloblastoma tumor samples, separated by inferred molecular subtype of medulloblastoma tumor samples. Crossbars indicate mean values.

**Supp. Fig. 12. Attempted inference of molecular subtype for glioma tumor samples**.

(a) PCA of glioma tumor samples using the top 5000 most variable genes by IQR across glioma tumor samples. (b) PCA of glioma tumor samples using 474 overlapping genes from the 500-gene set introduced in Teo et al. 2019. (c) Heatmap of glioma tumor samples using 474 overlapping genes from the 500-gene set introduced in Teo et al. 2019. (d) PCA of GBM tumor samples using 474 overlapping genes from the 500-gene set introduced in Teo et al. 2019. (e) Heatmap of GBM tumor samples using 474 overlapping genes from the 500-gene set introduced in Teo et al. 2019.

**Supp. Fig. 13. Attempted inference of molecular subtype for neuroblastoma tumor samples**.

(a) PCA of neuroblastoma tumor samples using the top 5000 most variable genes by IQR across neuroblastoma tumor samples. (b) Heatmap of neuroblastoma tumor samples using the top 5000 most variable genes by IQR. (c) PCA of neuroblastoma tumor samples using 1,248 overlapping super enhancer target genes from Gartlgruber et al. 2020. (d) Heatmap of neuroblastoma tumor samples using 1,248 overlapping super enhancer target genes from Gartlgruber et al. 2020.

