## Supplemental Figures for "Transcriptional fidelity enhances cancer cell line selection in pediatric cancers"

### Supplemental Figure 1

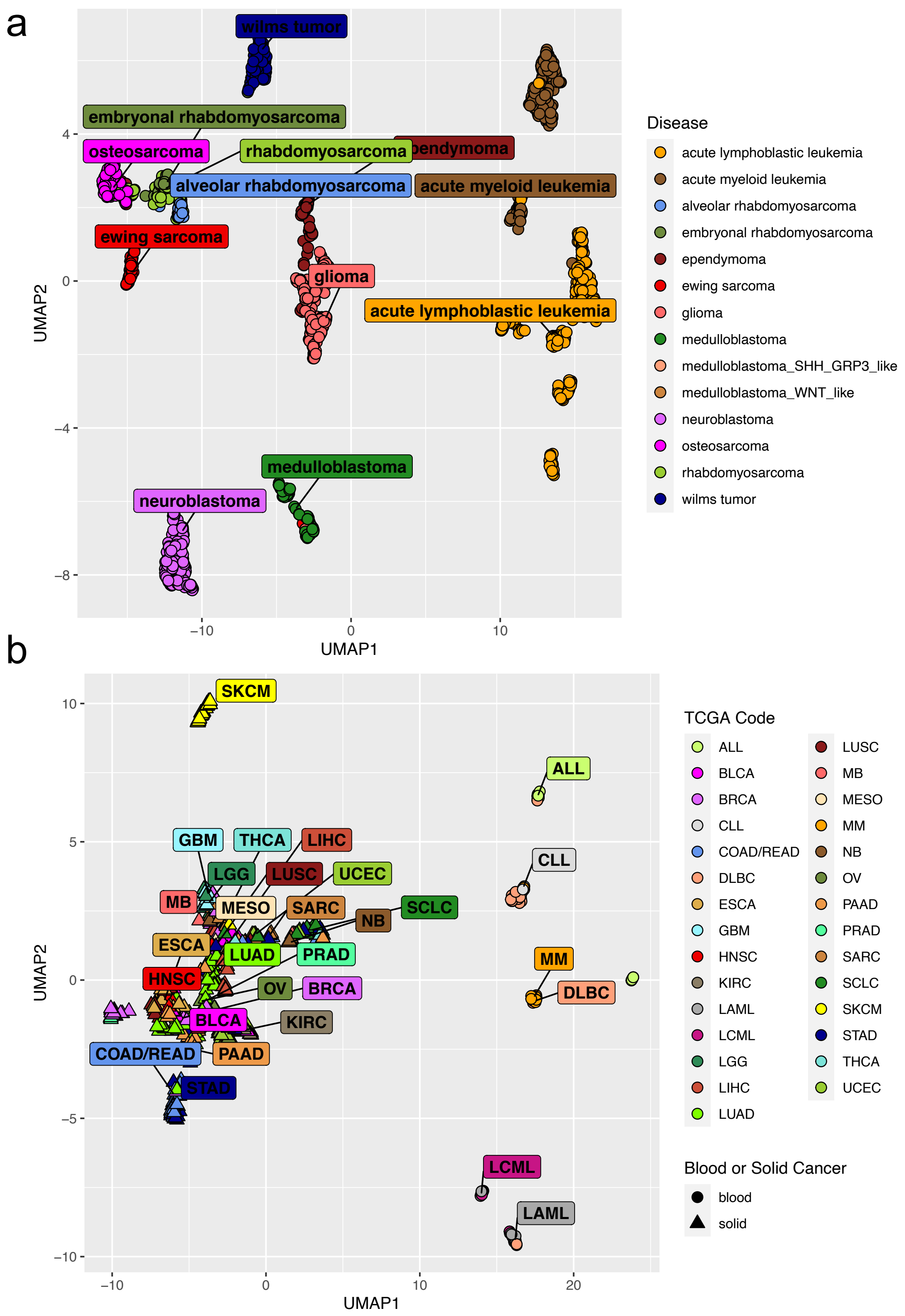

### Supplemental Figure 2

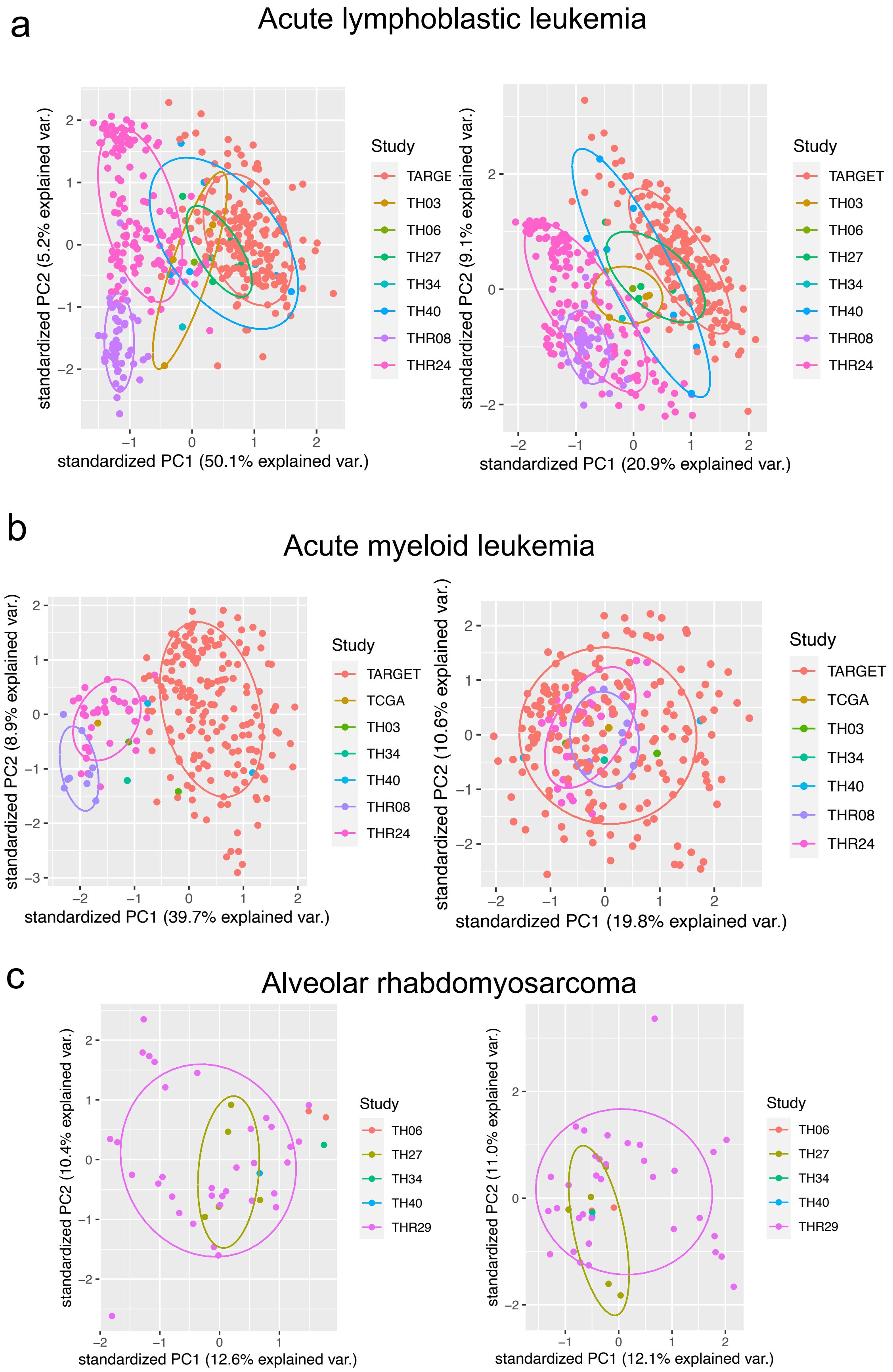

Supplemental Figure 3

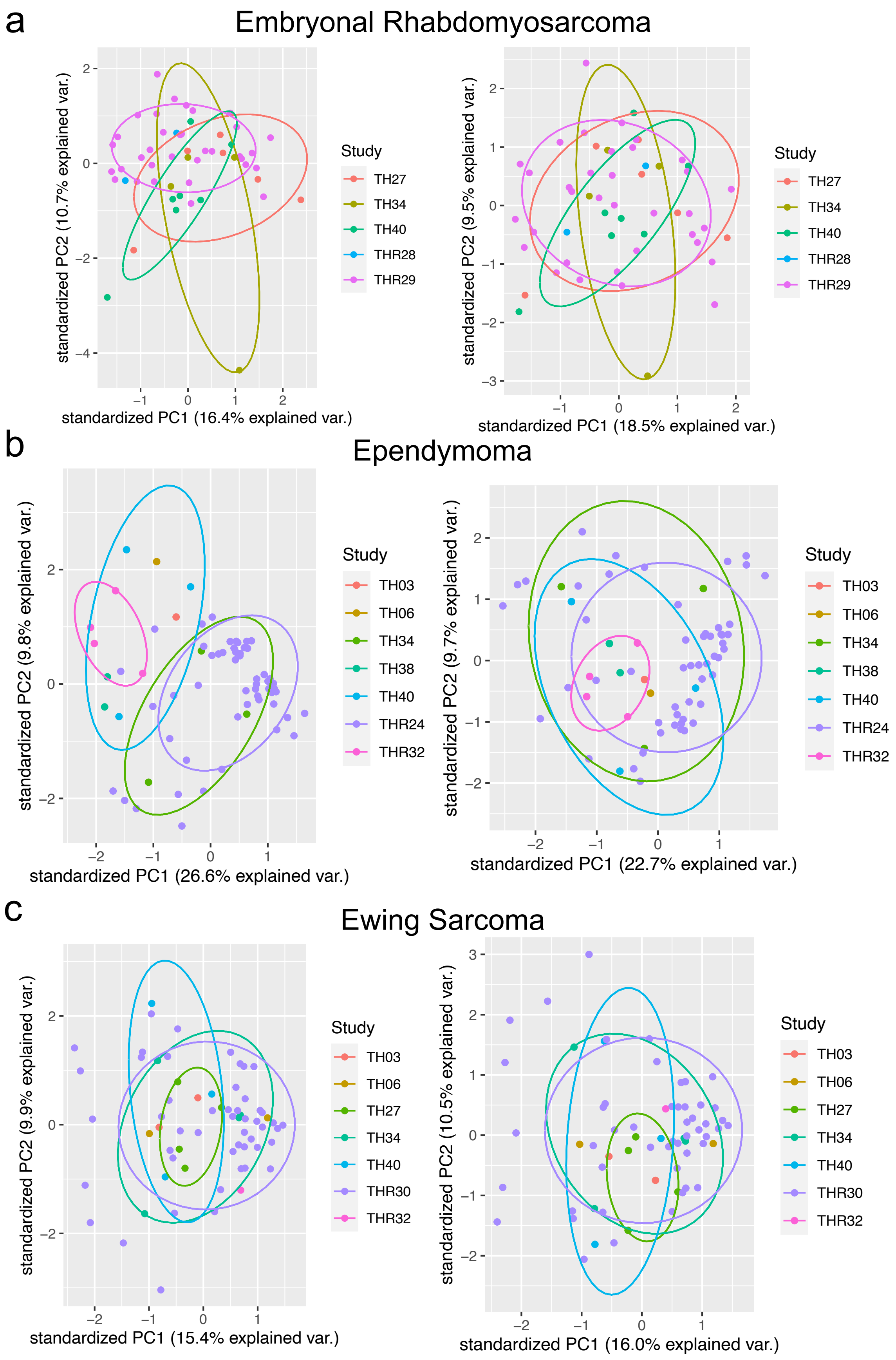

### Supplemental Figure 4

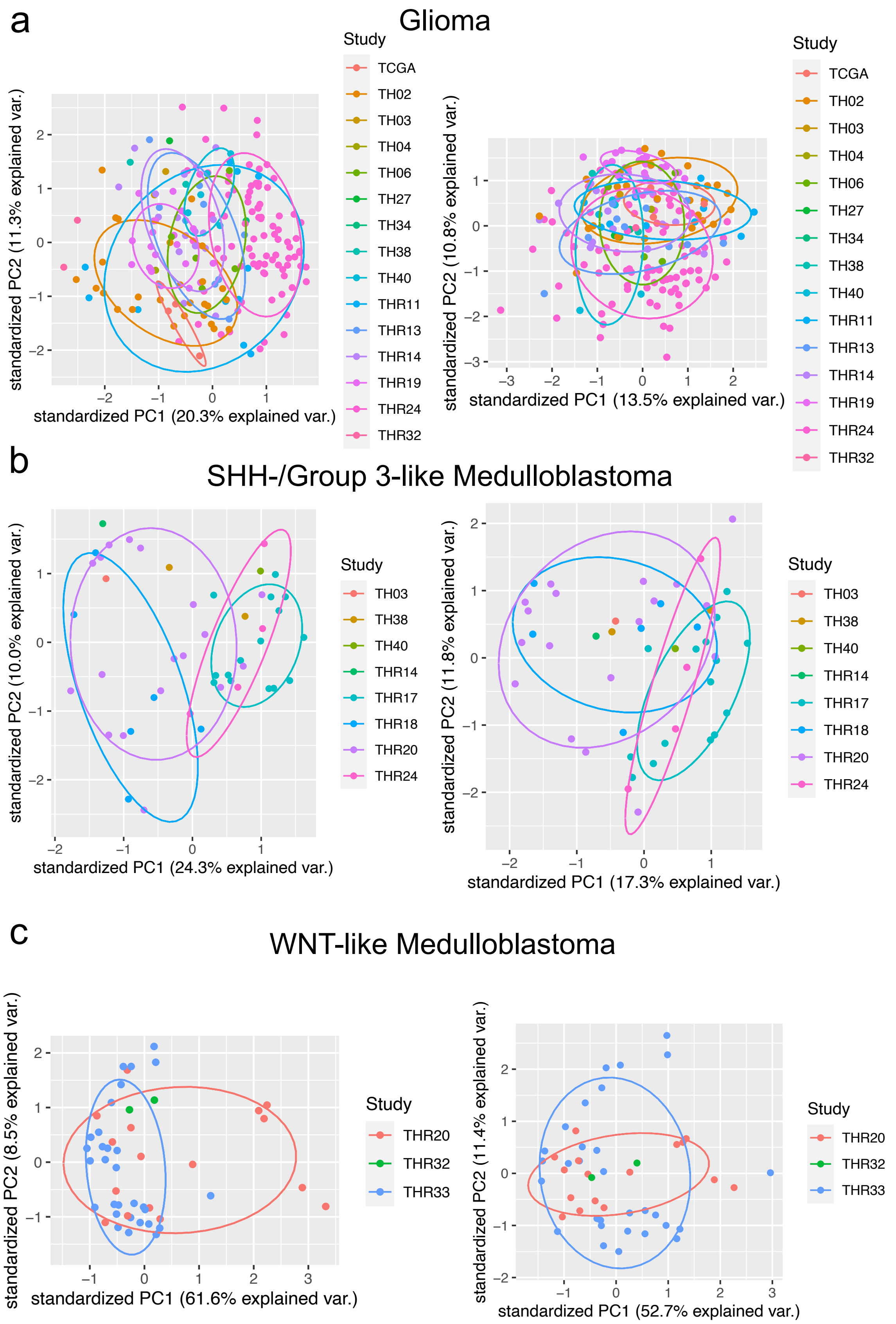

### Supplemental Figure 5

a

#### Neuroblastoma

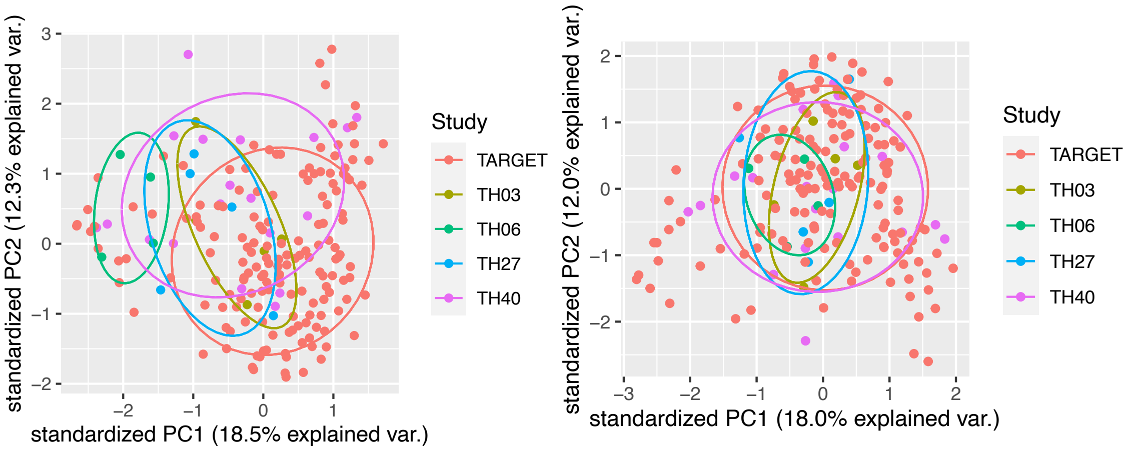


b

#### Osteosarcoma

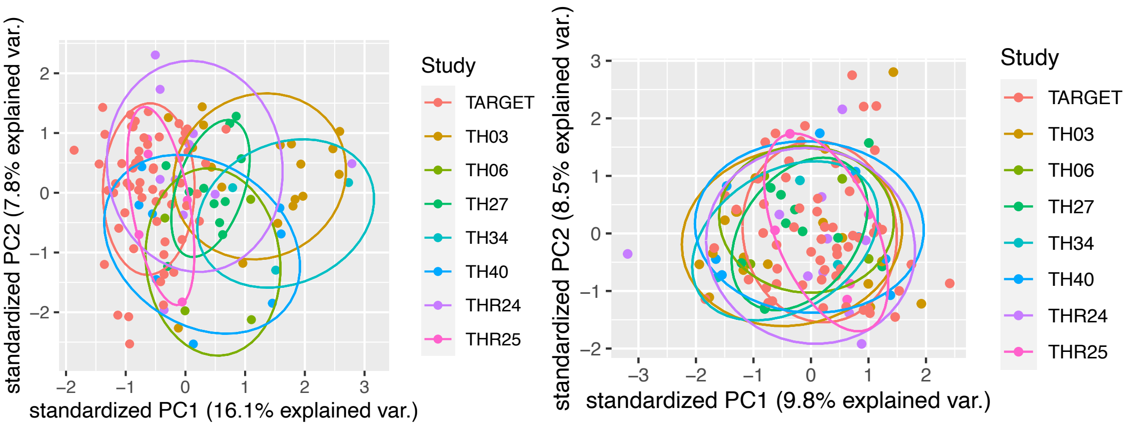


c

#### Rhabdomyosarcoma

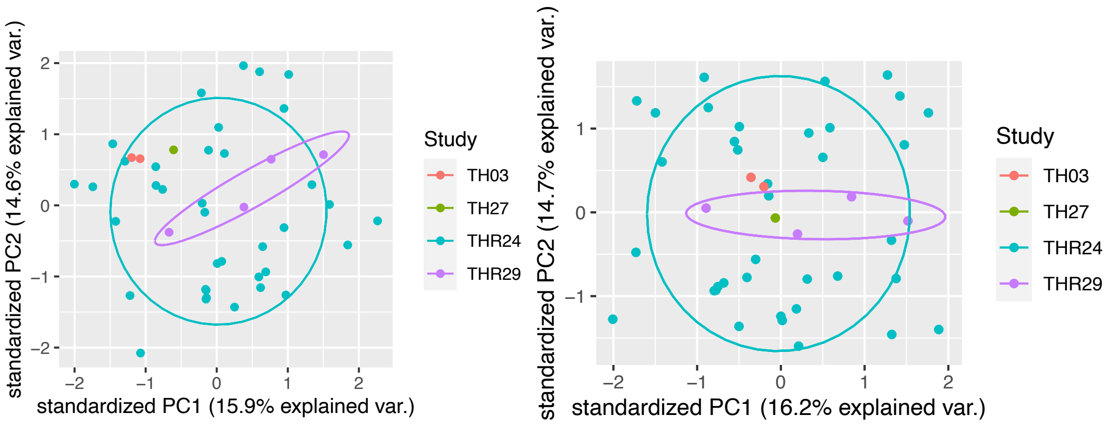

### Supplemental Figure 6

Wilms Tumor

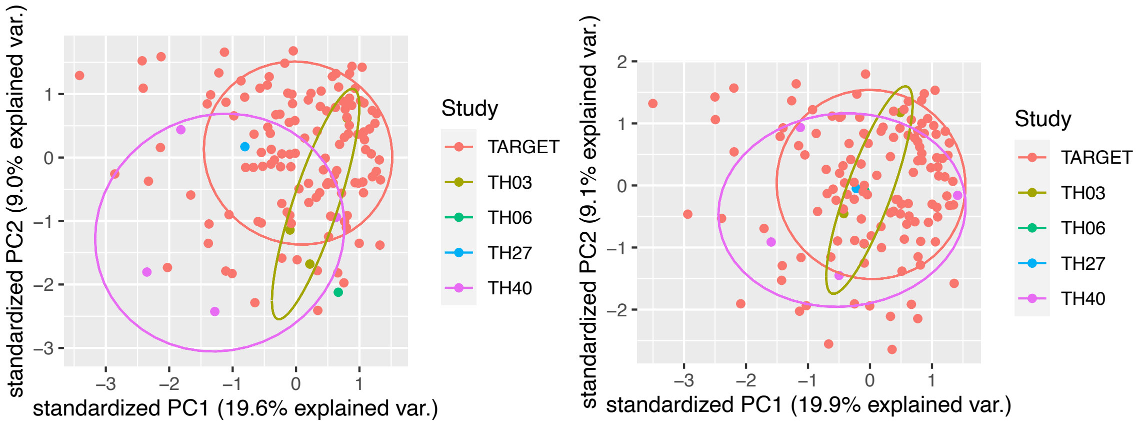

### Supplemental Figure 7

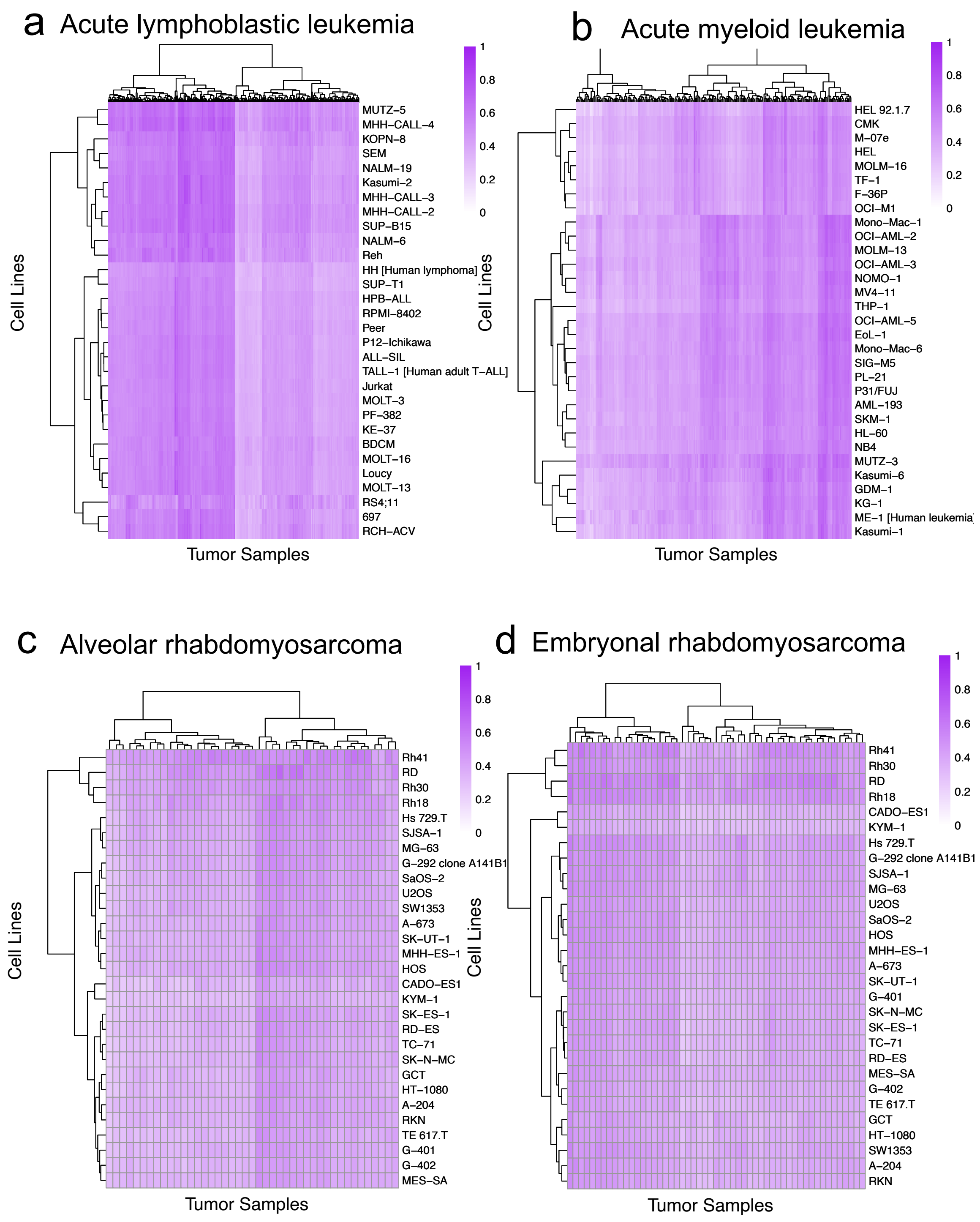

### Supplemental Figure 8

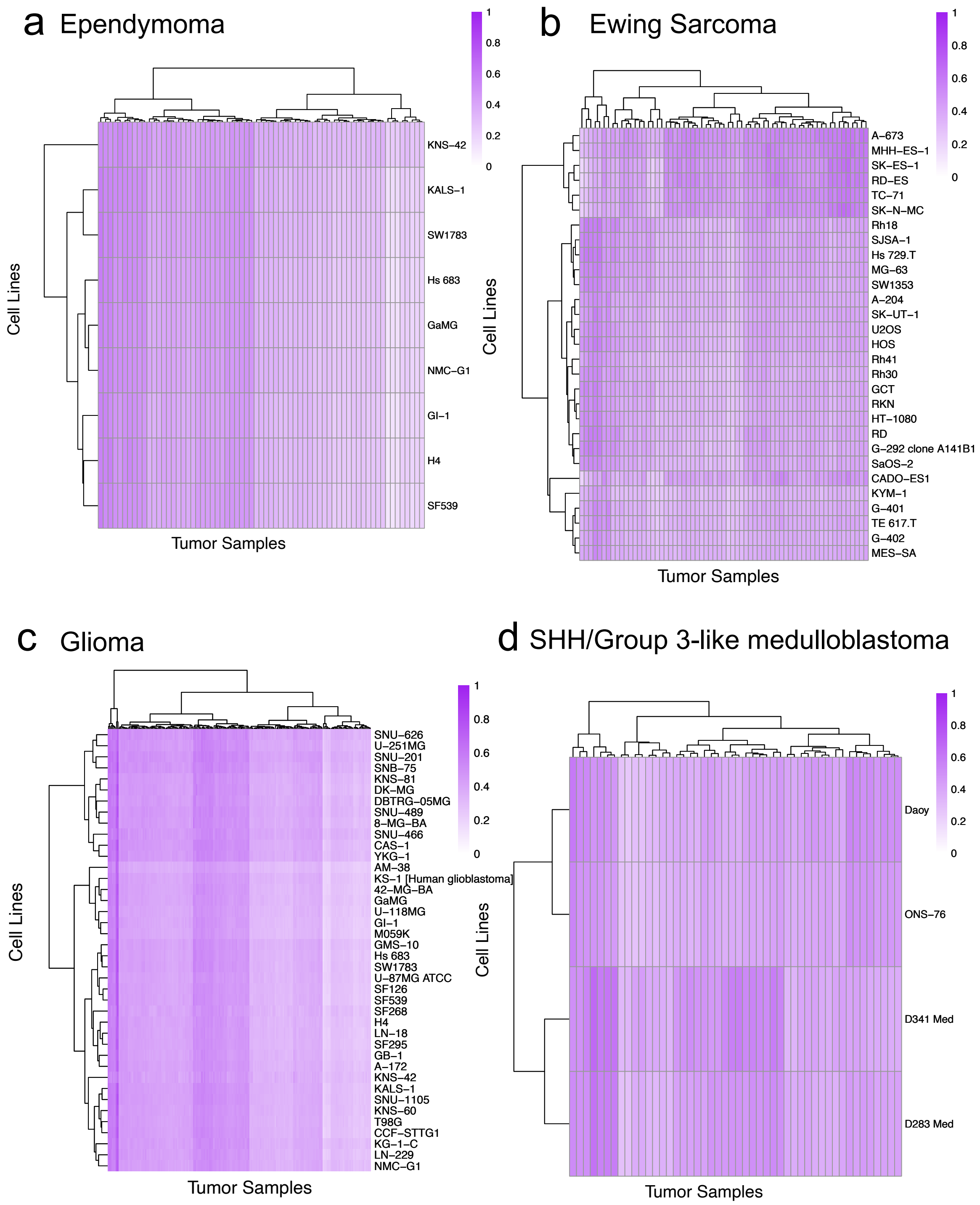

Supplemental Figure 9

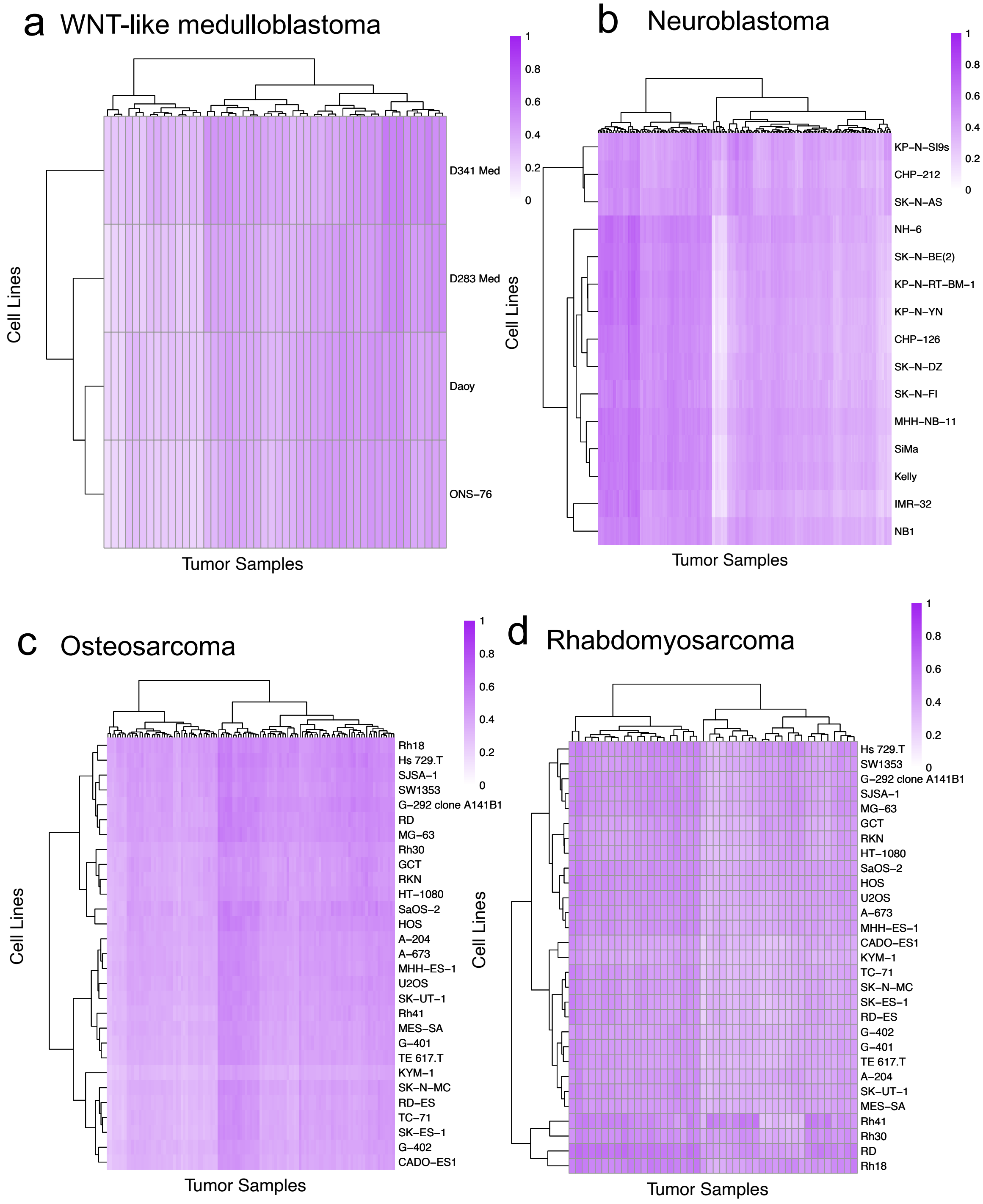

### Supplemental Figure 10

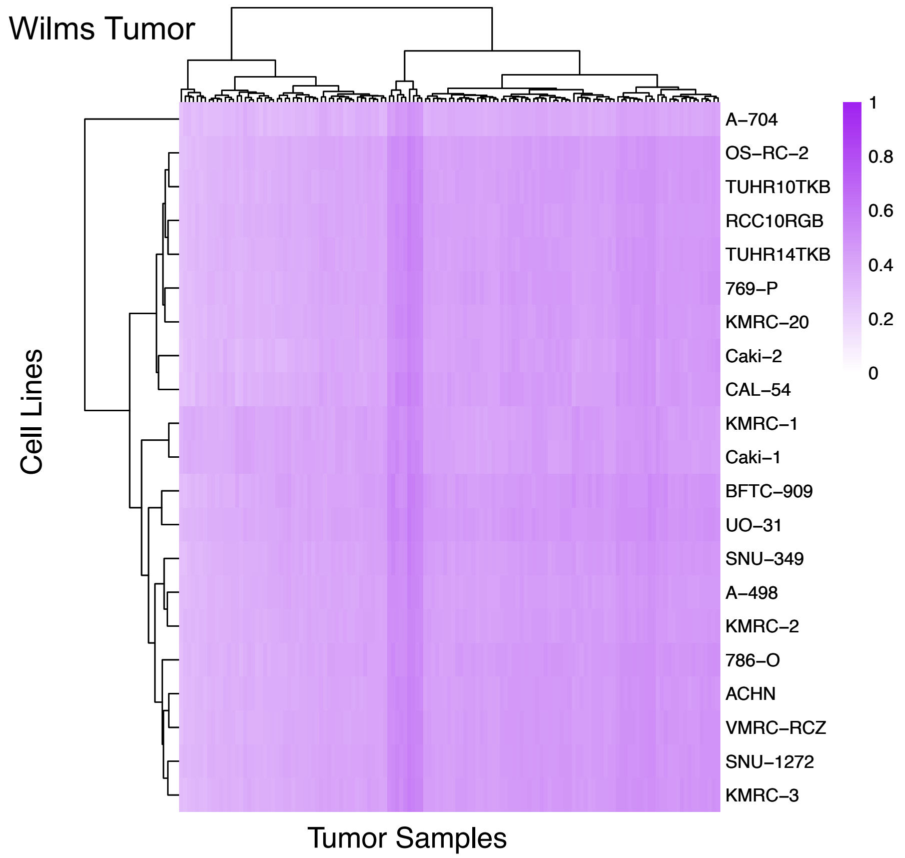

Supplemental Figure 11

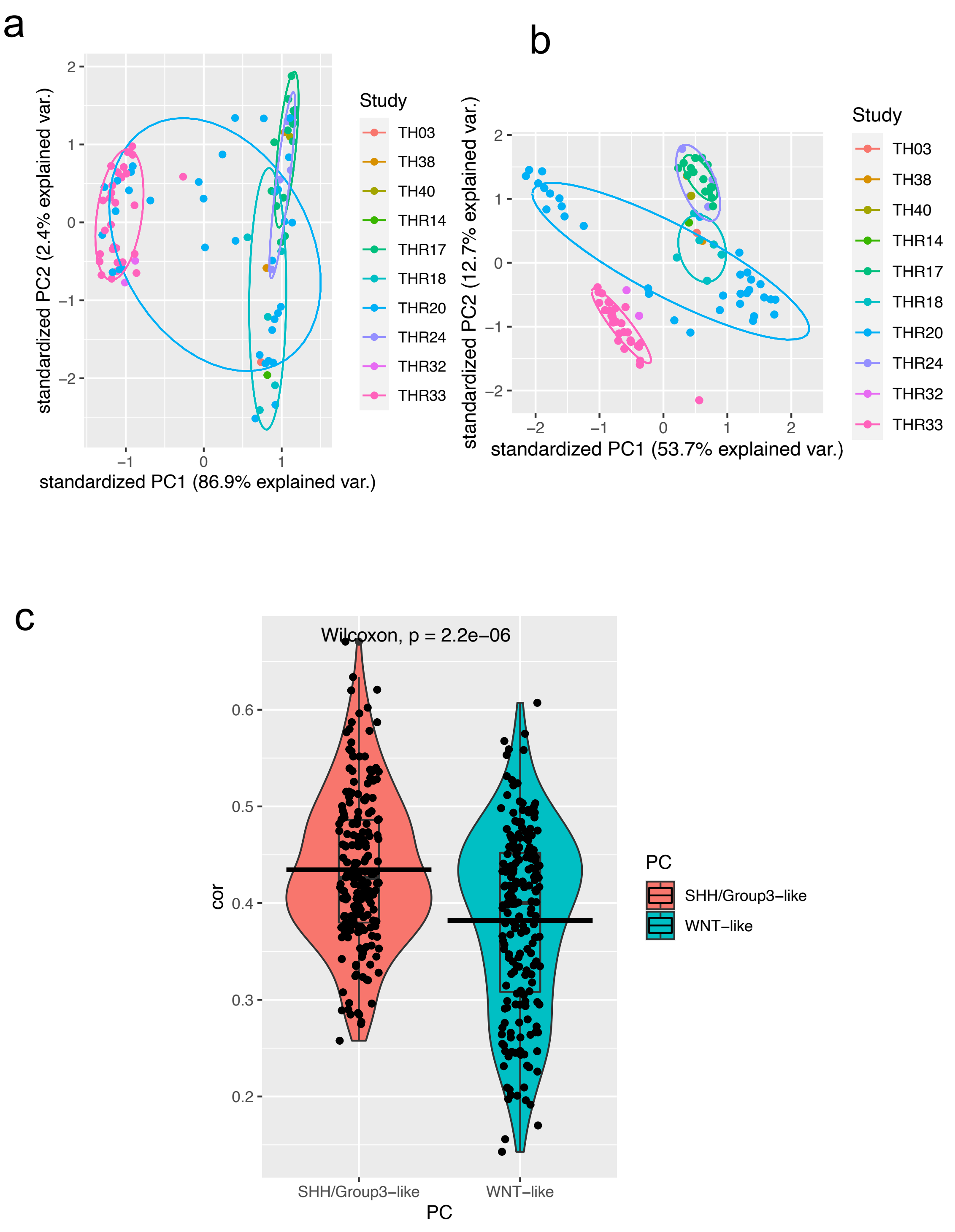

### Supplemental Figure 12

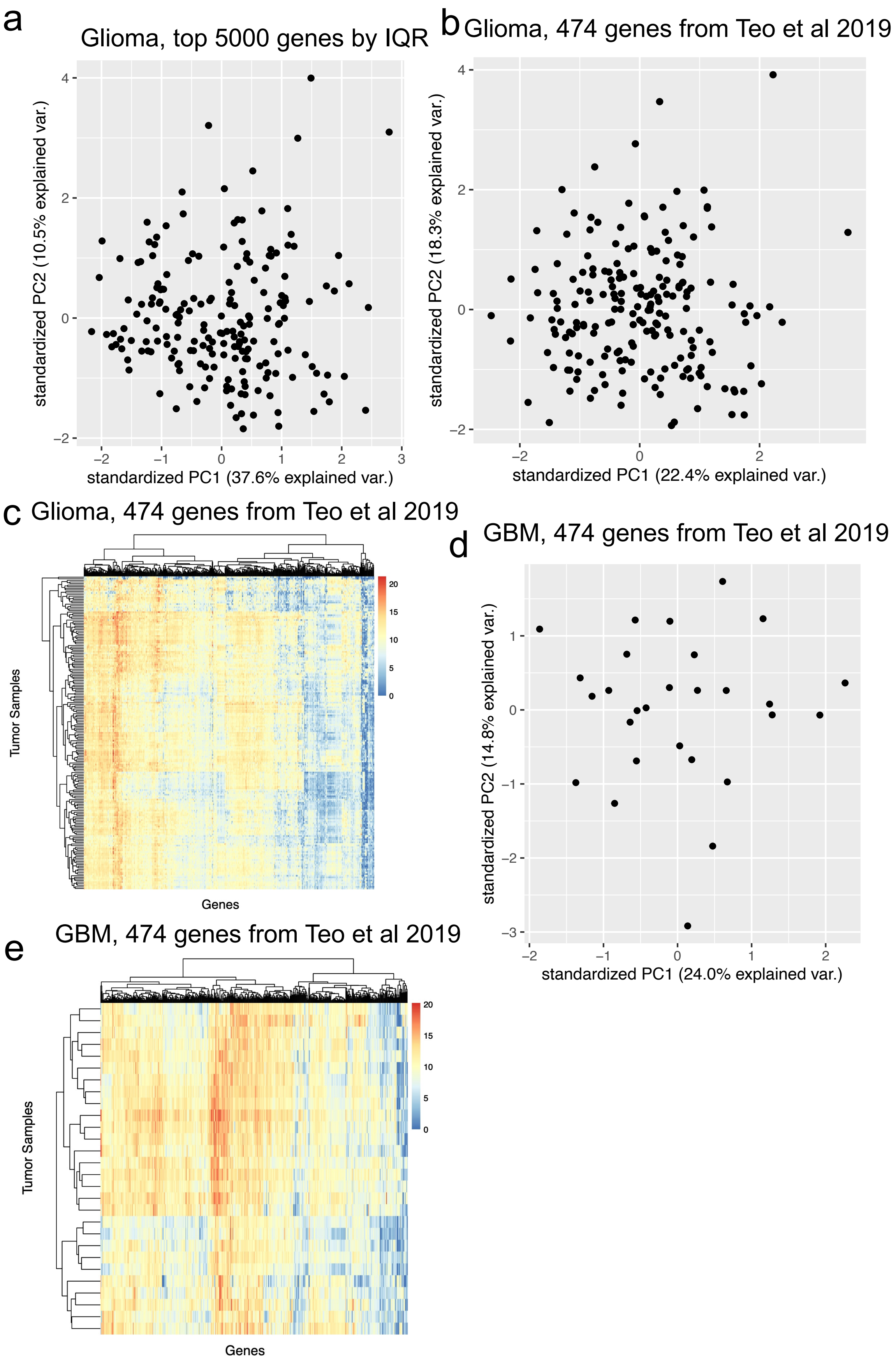

Supplemental Figure 13

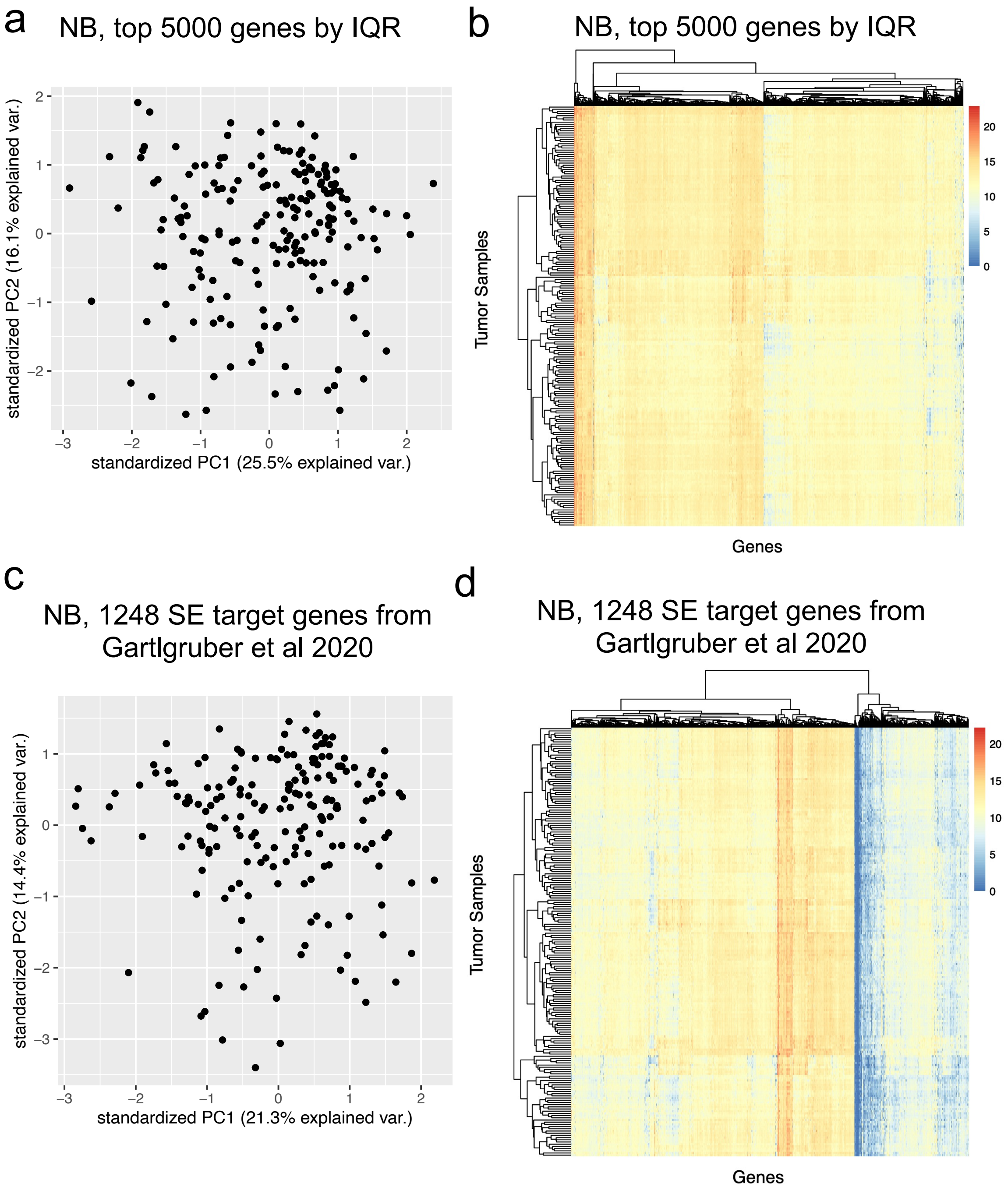
